# Machine learning-assisted fluoroscopy of bladder function in awake mice

**DOI:** 10.1101/2022.04.12.488006

**Authors:** Helene De Bruyn, Nikky Corthout, Sebastian Munck, Wouter Everaerts, Thomas Voets

## Abstract

Understanding the lower urinary tract (LUT) and development of highly needed novel therapies to treat LUT disorders depends on accurate techniques to monitor LUT (dys)function in preclinical models. We recently developed videocystometry in rodents, which combines intravesical pressure measurements with X-ray-based fluoroscopy of the LUT, allowing the *in vivo* analysis of the process of urine storage and voiding with unprecedented detail. Videocystometry relies on the precise contrast-based determination of the bladder volume at high temporal resolution, which can readily be achieved in anesthetized or otherwise motion-restricted mice but not in awake and freely moving animals. To overcome this limitation, we developed a machine-learning method, in which we trained a neural network to automatically detect the bladder in fluoroscopic images, allowing the automatic analysis of bladder filling and voiding cycles based on large sets of time-lapse fluoroscopic images (>3 hours at 30 images/second) from behaving mice and in a non-invasive manner. With this approach, we found that urethane, an injectable anesthetic that is commonly used in preclinical urological research, has a profound, dose-dependent effect on urethral relaxation and voiding duration and that implantation of a suprapubic catheter, as is standardly performed for cystometric analyses, leads to a ∼4-fold reduction in bladder capacity. Our findings provide a paradigm for the non-invasive, *in vivo* monitoring of a hollow organ in behaving animals and pinpoint important limitations of the current gold standard techniques to study the LUT in mice.

## Introduction

The lower urinary tract (LUT), consisting of the bladder and urethra, plays an important role in our daily life functioning. It is responsible for storing urine and emptying the bladder at an appropriate time and place. Proper LUT functioning is the result of complex communication pathways between the lower urinary tract and central nervous system (CNS). An estimated 20% of the population suffers from some sort of lower urinary tract dysfunction (LUTd), such as urinary incontinence, frequency, urgency, or bladder pain, which can have a strongly negative impact on health and wellbeing. Current treatments for LUTd generally lack efficacy and often come with bothersome side effects. Therefore, there is a high need for new treatment options, which depend on adequate preclinical models to study mechanisms of LUT control and to test potential novel therapies.

Due to their anatomical and physiological similarities to humans, their cost-effectiveness, and the availability of a vast number of genetically modified strains, mice are favored models in preclinical LUT research (1). In the last decades, cystometry has been considered the gold standard technique to study bladder function in mice *in vivo*. Cystometry is an experimental procedure whereby the bladder is continuously filled via an implanted catheter, and pressure inside the bladder is monitored during multiple filling and voiding cycles. More recently, this technique has sometimes been complemented with urethral electromyography (UEM) to provide simultaneous information regarding the activity of the urethral sphincters (2). Whereas these approaches have fueled important advances in our understanding of the cellular and molecular mechanisms that steer the LUT, they also come with important drawbacks. First, cystometry and UEM do not provide quantitative information regarding the filling state of the bladder or the urethral flow, whereas changes in these parameters are hallmarks of various LUTds. Second, cystometry and UEM require the introduction of respectively a catheter in the bladder wall and metal electrodes in the urethral sphincter, both of which can affect LUT function. Third, cystometry and UEM are mostly performed on urethane-anesthetized or physically restrained mice to avoid complications and artifacts due to animal movement. The consequences of these procedures on LUT function in mice are incompletely understood.

To overcome some of these limitations, we recently introduced videocystometry, a new approach combining cystometry with continuous, fluoroscopy-based bladder imaging. By imaging the contrast-filled bladder at video rate, this approach provides quantitative and time-resolved measurements of bladder volume and urethral flow, along with other relevant aspects of LUT function such as vesicoureteral reflux and micromotions of the bladder wall (3). Notably, the accurate analysis of bladder volume and urethral flow during consecutive filling-voiding cycles relies on the precise delineation of the bladder. This is readily achieved in anesthetized or otherwise motion-restrained mice, where the constant background of fluoroscopic images can be subtracted. However, in awake, unrestrained, and behaving animals, the position of the bladder relative to other structures visible in the fluoroscopic images (e.g. bones) changes constantly. Therefore, our earlier analyses relied on manual tracking of the border of the bladder, which is feasible for a limited number of fluoroscopic images but clearly unworkable for typical minutes-to-hours-long imaging sessions at 30 images per second, yielding >10^5^ images.

To address this problem, we developed a machine-learning approach where we trained a neural network that automatically and faithfully detects the bladder in large sets of time-lapse fluoroscopic images in mice, allowing accurate determination of volume- and flow-related parameters in behaving mice. The combination of videocystometry and machine learning allowed us to pinpoint the profound effects of urethane, an injection anesthetic standardly used in preclinical bladder research, on the function of the LUT. Furthermore, we provide proof of principle for the use of fluoroscopy combined with machine-learning-based image analysis to non-invasively monitor bladder function in awake animals, which reveals the significant impact of catheterization on functional bladder capacity.

## Results

### Machine-learning-based analysis of fluoroscopic images of the LUT

We combined classic cystometry with fluoroscopy-based imaging to study LUT function in awake, unrestrained mice. In previous work, we had established two methods to determine bladder volume (V_ves_) from fluoroscopic images. The first method, which is based on the linear relation between background-corrected opacity and volume of contrast-filled bladders, is only applicable to recordings with a stable background, i.e. in anesthetized or otherwise immobile animals. The second method, which is based on the approximation of the bladder as a spheroid, depends on the delineation of the bladder wall but does not require background subtraction, making it, in principle, suitable to determine bladder volume from images of moving mice. However, considering that our videocystometry experiments lasted up to 3 hours, at a sampling rate of 30 images/s, manual delineation of the bladder wall in these large sets of time-lapse fluoroscopic images was impossible. We therefore developed a machine-learning protocol to automatically identify the bladder border in fluoroscopy images, independent of the position or posture of the mouse or the degree of bladder filling. To achieve this, we performed several cycles of training of a neural network using a set of human-annotated fluoroscopy images (Fig. 1a).

**Figure 1:**
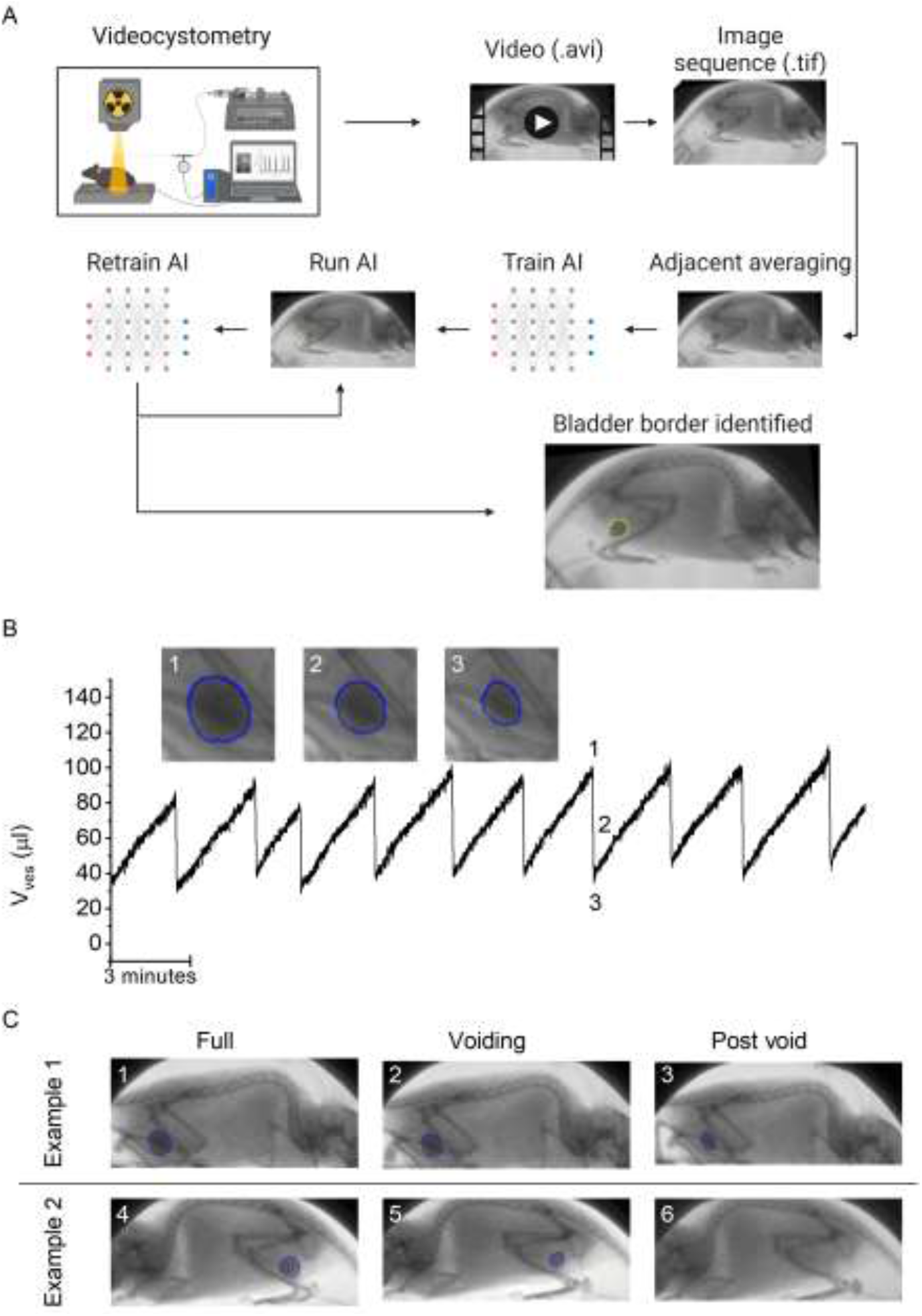
A machine learning protocol for the automated annotation of the bladder in fluoroscopic images. **(A)** Image analysis protocol using artificial intelligence-based automatic annotation of the bladder border. **(B)** Representative bladder volume trace based on artificial assisted automated annotation of the bladder. Annotated bladder images (blue line), corresponding to different filling states of the bladder, are shown. **(C)** Examples showing annotated bladders before, during and after voiding in different positions.

The machine-learning protocol was able to successfully identify the bladder border in time-lapse image sequences, irrespective of the posture of the animal or phase of the filling/voiding cycle (fig. 1b-c), with the exception of brief failures of one or two frames during abrupt mouse movements or when the bladder was fully emptied. This approach allowed us to determine time-dependent changes in bladder volume based on the spheroid approximation (fig. 1b), along with other volume-related voiding parameters: bladder capacity (BC), residual volume (RV), and voiding efficiency (E_voiding_), as described previously (3).

### Dose-dependent effect of urethane anesthesia on voiding function

Next, we used this approach to evaluate the dose-dependent effects of urethane, the most predominantly used anesthetic during *in vivo* cystometry in rodents, on LUT function. Urethane was administered at 4 time-points, leading to a stepwise increase of the total dose from zero to 1.2 g/kg bodyweight, the latter being the typical dose used in most rodent studies. This protocol caused the animals to gradually transit from a non-anesthetized state, where they could freely move within the boundaries of the recording box, to a fully anesthetized, largely immobile state. Before dosing and after each urethane administration, videocystometry was performed during 40 minutes (n=5), resulting in 200-min-long recordings. These encompass approximately 360,000 time-lapse fluoroscopy images per experiment, from which we derived time-dependent changes in volume and the various volume-related parameters (Fig. 2a). Urethane did not have a statistically significant effect on BC (Fig. 2c), but exhibited a pronounced and dose-dependent effect on RV and E_voiding_ (fig. 2 d, e). Indeed, whereas awake animals completely emptied their bladder at each void, urethane at doses >0.3g/kg caused a dose-dependent increase in the volume that remained after each void and a corresponding decrease in E_voiding_ from ∼100% to below 50% (fig. 2e). The intercontractile interval (ICI) decreased concomitantly with the reduced efficiency of voiding (fig. 2f). Note that animals that were immediately treated with a single 1.2g/kg-dose yielded similar values for BC, RV and E_voiding_ as animals that reached this dose at the end of the stepwise dosing protocol, indicating that the reduced voiding efficiency and increased residual volume are due to the urethane rather than to fatigue (fig. 2c-f, blue) (3). Taken together, our findings indicate that urethane leads to a pronounced and dose-dependent reduction of the ability of the bladder to empty during voiding.

**Figure 2:**
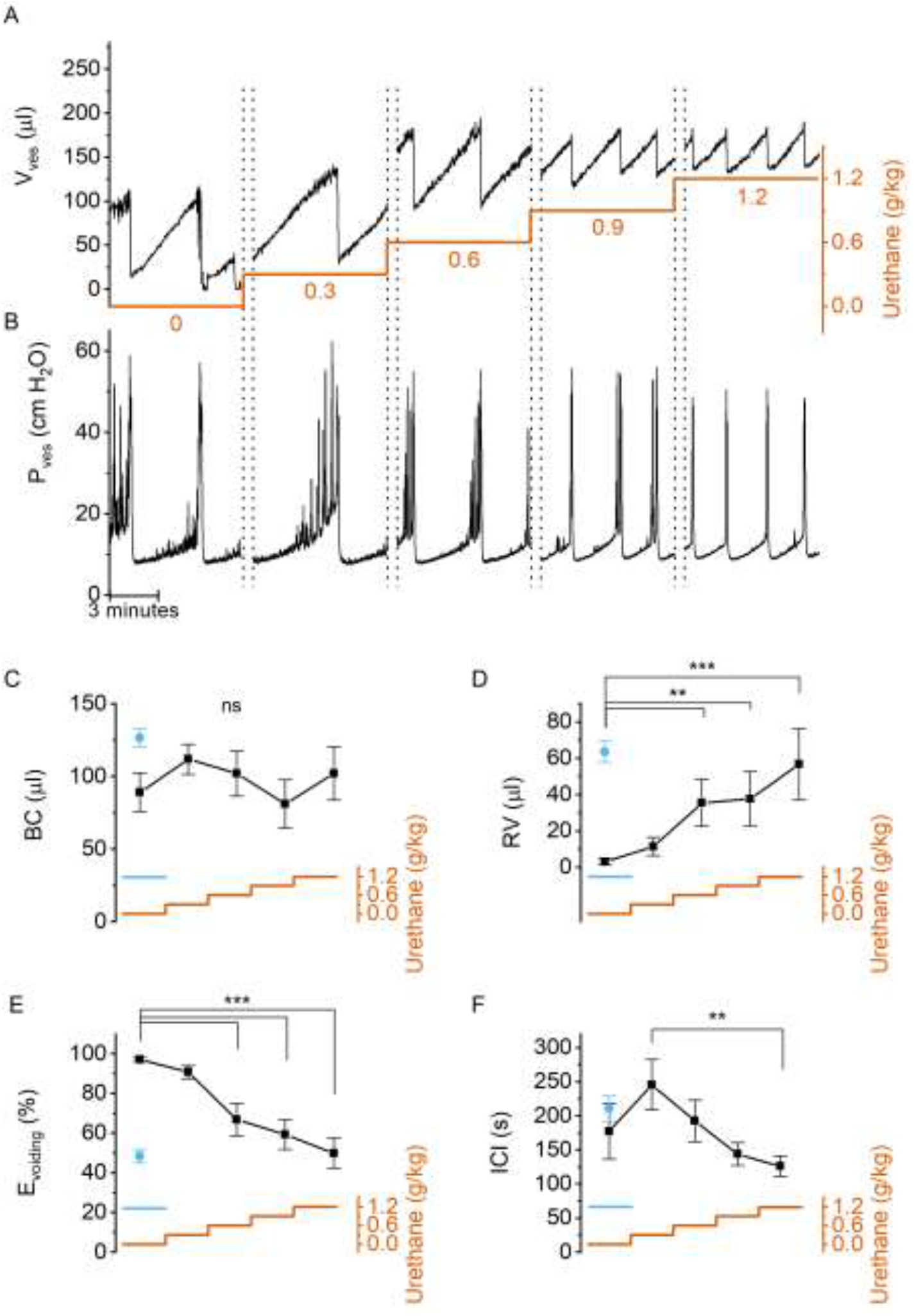
Dose-dependent effects of urethane on bladder volume and voiding efficiency. **(A**,**B)** Short excerpts of simultaneous bladder volume and intravesical pressure recordings traces at increasing cumulative doses of urethane. **(C-F)** Average bladder capacity, residual volume, voiding efficiency and intercontractile interval at increasing doses of urethane (mean ± SEM). For comparison. The blue symbols represent data from mice that received a single dose of 1.2 g/kg at the start of the recording (3). One-way repeated measures ANOVA was used to test for statistically significant differences with the 0.0 g/kg condition. **, ***: p<0.01, 0.001.

Next, we zoomed in on individual voids to identify changes in void duration or urethral flow that could explain the dose-dependent reduction in voiding efficiency by urethane (fig. 3a). Increasing doses of urethane resulted in a gradual decrease in void duration, as quantified by the 20-80% time interval (t_20-80_) (fig. 3c). Urethane also affected the urethral flow rate (UFR), which we determined from the time derivative of the changes in bladder volume, showing a gradual decrease at increasing doses (fig. 3d). These findings demonstrate that incomplete bladder emptying at higher urethane doses is the result of both shorter void duration and reduced flow rate of urine expulsion from the bladder during a void. Analysis of the combined intravesical pressure and UFR measurements further revealed that the urethane-induced reduction of urethral flow was the result of a lower urethral flow conductance (UFC), while intravesical pressure during a void was not significantly affected (fig. 3e). Taken together, these findings reveal that urethane has a profound and dose-dependent effect on voiding efficiency, which can be attributed to a reduced urethral relaxation and shorter void duration. Moreover, when we analyzed the 80-s period preceding each void, we found a dose-dependent decrease in the number of non-voiding contractions (fig 3b, f).

**Figure 3:**
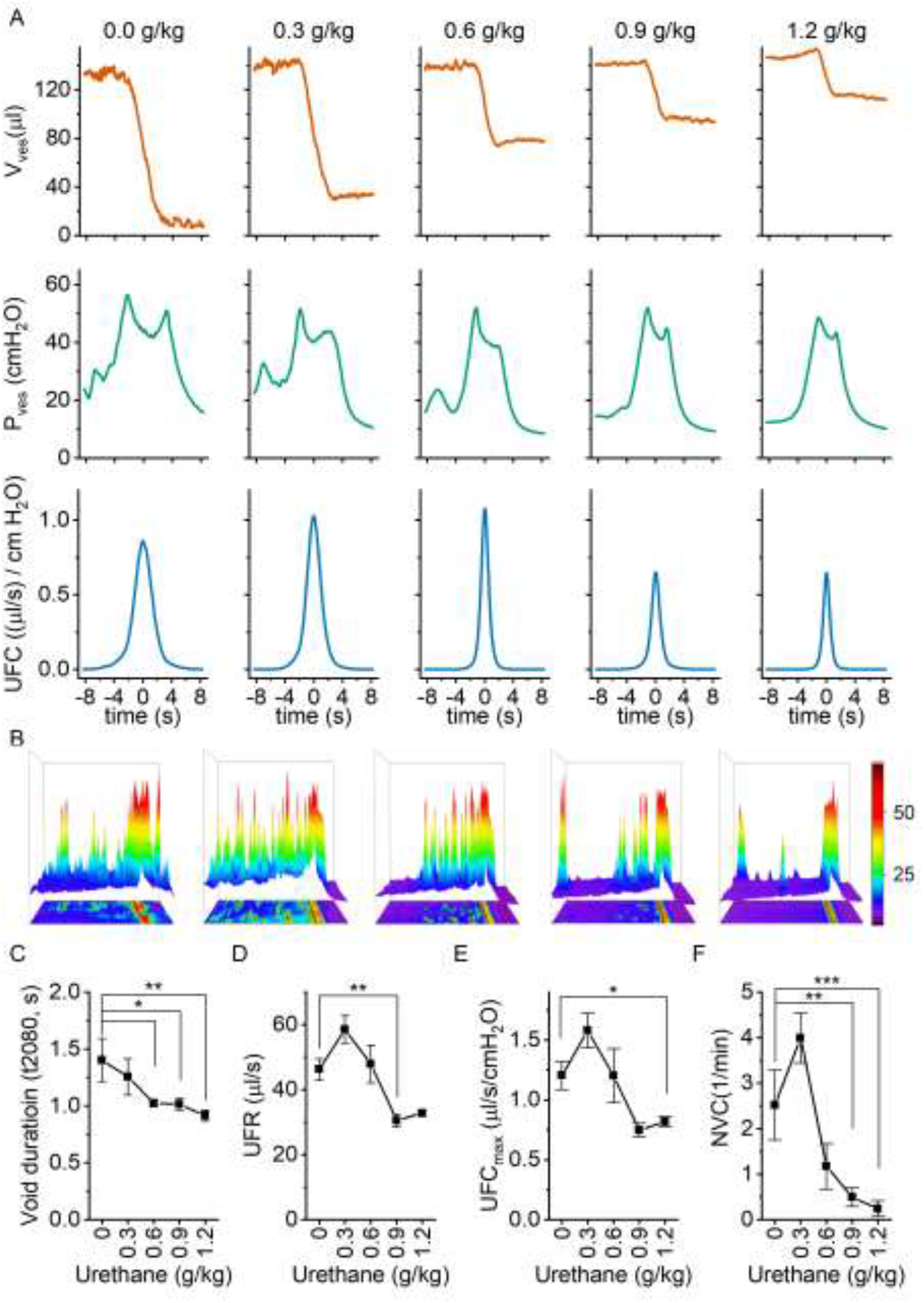
Dose-dependent effects of urethane on the voiding process. **(A)** Zoomed-in examples of average bladder volume, bladder pressure and urethral flow conductance at increasing doses of urethane. **(B)** Combined 2D/3D pressure plots showing 100 s of intravesical pressure changes aligned to a voiding contraction at time-point 80s from a single animal at the indicated dose. Each row represents the pressure during one individual void. The color scale on the right represents the intravesical pressure (in cmH_2_O). **(C-F)** Assessment of the effect of urethane on void duration, urethral flow rate, urethral flow conductance and non-voiding contractions (mean ± SEM). One-way repeated measures ANOVA was used to test for statistically significant differences with the 0.0 g/kg condition. *,**, ***: p<0.05, 0.01, 0.001.

### Non-invasive fluoroscopic monitoring of the LUT

In classic cystometry and videocystometry experiments, surgery is performed to implant a catheter into the bladder dome, through which the bladder is filled at a constant, often supraphysiological filling rate. The impact of this invasive intervention on bladder function is poorly understood. We adapted our fluoroscopy-based approach to monitoring bladder volume in a non-invasive manner in awake, unrestrained animals. Injection of a bolus of contrast solution subcutaneously in the scruff region resulted in the excretion of iodine-containing urine, enabling fluoroscopy-based imaging of the bladder without catheter implantation. In this setting, the filling rate of the bladder is solely determined by the renal urine output, which we accelerated within a physiologically acceptable range using a diuretic. Animals were imaged during 120 minutes, and the machine-learning protocol was used to automatically detect the bladder border. This approach, fluoroscopic volumetry, allowed for the first time to accurately and continuously quantify volume- and flow-related parameters non-invasively in awake and behaving animals (fig. 4a-c), albeit without concomitant pressure measurement.

**Figure 4:**
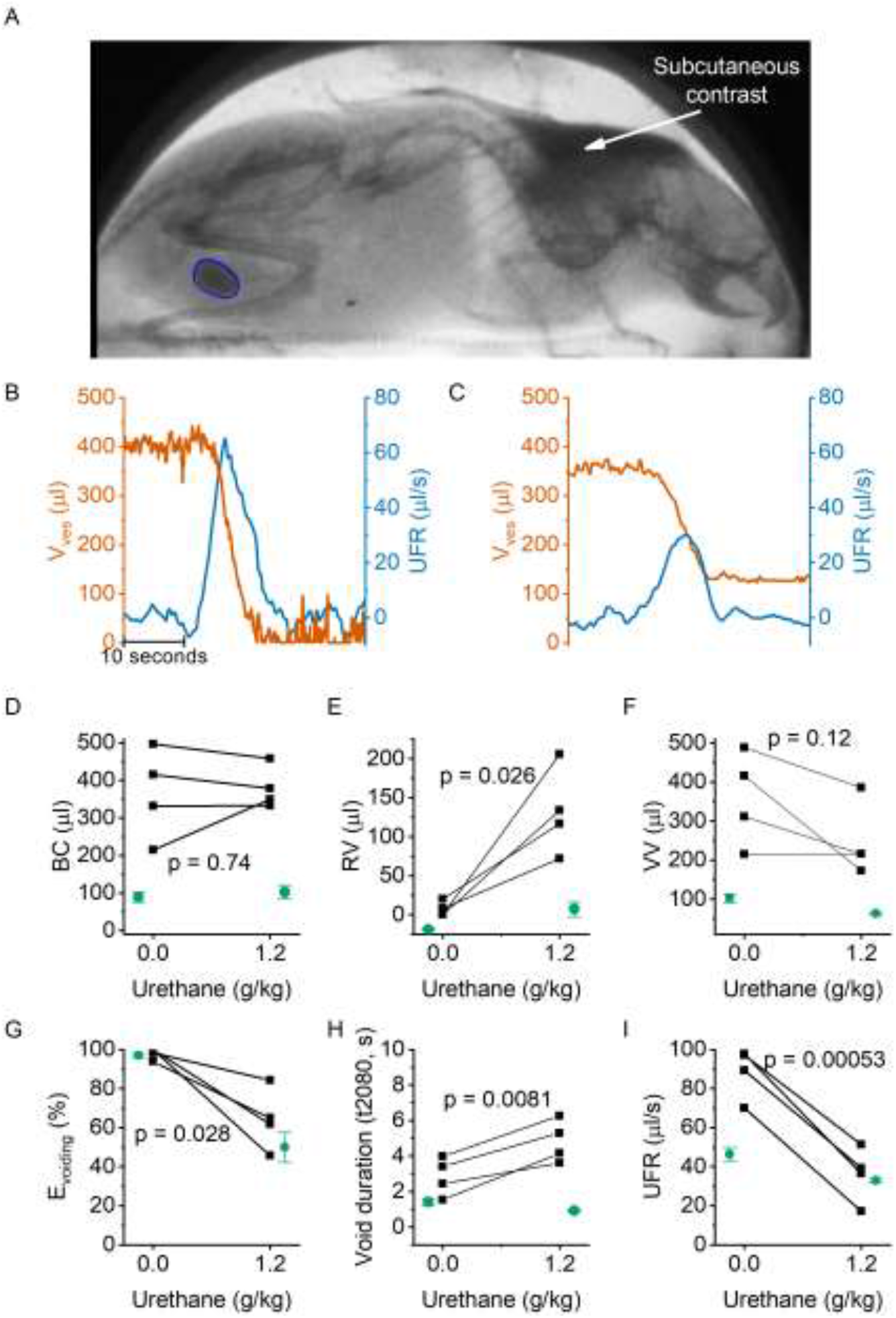
Non-invasive fluoroscopic volumetry. **(A)** Fluoroscopic image showing the presence of a contrast agent in the scruff (arrow) and in the bladder of an awake and non-restrained mouse, as well as the automatic annotation of the bladder border. **(B-C)** Zoomed-in example of bladder volume and urethral flow rate in an animal that did not receive any urethane anesthesia and in the same animal after dosing at 1.2 g/kg. Note that, due to the complete bladder emptying and slow filling in the absence of urethane, the void is followed by a period of low-accuracy volume measurement. **(D-I)** Comparison of the indicated voiding parameters in animals without an implanted catheter (black), before and after urethane dosing. In green, we show the corresponding values from catheterized animals reproduced from figures 2 and 3. Paired t-tests comparing the values in the non-catheterized animals before and after urethane dosing yielded the indicated p-values.

We performed fluoroscopic volumetry in four animals, first in the awake conditions and subsequently after administration of urethane at a dose of 1.2 g/kg. Notably, both in the awake and in the anesthetized conditions, we observed a dramatically larger BC in comparison with catheterized animals: BC amounted to 364 ± 60 µl (compared to 89 ± 13 µl in catheterized animals, p = 0.0015) in the absence of urethane, and to 380 ± 28 µl (compared to 102 ± 18 µl, p = 0.02) after dosing 1.2 g/kg urethane (fig. 4d). Like in the catheterized animals, we detected complete voiding in the awake animals, whereas anesthetized animals had substantial residual volume, corresponding to a voiding efficiency of ∼60% (fig. 4e-g). In comparison with catheterized animals, both the maximal UFR and the void duration were significantly increased (fig. 4h-i). Notably, whereas urethane caused a shortening of the void duration in catheterized animals (fig. 3), a prolongation was observed in the non-operated animals (fig. 4h). Taken together, these results demonstrate the feasibility of machine learning-assisted bladder fluoroscopy to study bladder function in a non-invasive manner and highlight important effects of catheter implantation on bladder filling and voiding.

## Discussion

The process of bladder filling and voiding is regulated by complex and incompletely understood signaling pathways between the central nervous system and the LUT, which coordinate the contractile state of the bladder muscle and urinary sphincters. Dysregulation of these processes can lead to a plethora of bothersome LUT disorders, for which current treatments are often unsatisfactory. Rodents are widely used in preclinical research aimed at understanding the (patho)physiology of the LUT and at the development of novel treatments. However, the current state-of-the-art methods and approaches to study bladder function in rodents have various drawbacks, which may affect interpretation and compromise translation of the findings to the human situation. For instance, the gold standard technique in the assessment of LUT function in rodents, cystometry, requires the chirurgical implantation of a catheter through the bladder wall - the consequences of this invasive technique on bladder sensitivity and storage capacity are poorly understood. Moreover, invasive techniques such as cystometry and UEM are generally performed on anesthetized or physically restrained mice, which may substantially affect LUT function. Existing non-invasive techniques in awake, freely moving rodents, which depend on the detection of voided urine on filter paper or balances, lack detailed temporal resolution and deliver only limited information on the voiding process. Finally, none of these techniques provides precise and continuous measurements of the bladder volume.

Recently, we introduced videocystometry, combining cystometry with high-speed fluoroscopy imaging of the contrast-filled bladder. We established that bladder volume and urethral flow can be accurately derived from fluoroscopic images, based either on image opacity after background correction or on precise annotation of the bladder border. In awake and non-restrained animals, determination of bladder volume based on image opacity is not feasible due to the continuously changing background of the behaving animal. Manual annotation of the bladder border is unworkable for typical hours-long imaging sessions at 30 images per second, yielding several >300,000 images per experiment. In this study, we have overcome this limitation by developing and successfully implementing a machine-learning protocol, which allowed the automated identification of the bladder border and derived volumetric parameters from large sets of time-lapse images in awake, behaving mice.

Next, we used the machine learning-assisted fluoroscopy to quantify the effect of anesthesia on LUT function. Urethane, which can be administered subcutaneously, intraperitoneal or intravenously, is the most commonly used anesthetic in LUT research, preferred above other anesthetics that are known to suppress the micturition reflex or that create a less stable anesthesia (4,5). Despite its broad use in preclinical LUT research, the mechanism of action of urethane is not clearly understood, and the impact on LUT physiology remains poorly understood (5–7). We performed videocystometry in mice, where we started the recording in awake animals and monitored changes in bladder function upon stepwise increases in urethane dose up to a final 1.2 g/kg, which is a standard dose in LUT research. Urethane did not affect intravesical peak pressure during voiding, but dose-dependently suppressed non-voiding contractions, in line with earlier studies (8). In addition, urethane caused a strong and dose-dependent reduction of the voiding efficiency. Indeed, whereas in the awake animals, the bladder was completely emptied at every void, residual volume increased with mounting doses of urethane, causing voiding efficiency to decrease to around 50% at the highest dose. As these findings conform to previously published data on bladder volume and voiding efficiency in urethane-anesthetized mice, the suppressive effect of urethane on voiding cannot be attributed to gradual fatigue of the detrusor muscle in the course of the experiment (3). Detailed analysis of individual voids allowed us to attribute the reduction in voiding efficiency to both shorter void durations and a reduction in maximal UFC. These results indicate that urethane reduces the duration and extent of urethral relaxation during a void, leading to a pronounced reduction in voiding efficiency. Throughout these experiments, we observed that animals receiving the lowest dose of urethane (0.3 g/kg) were awake and behaving, but displayed a more tranquil behavior than non-anesthetized animals. Likewise, both pressure and volume data showed less artefacts in these animals treated with 0.3 g/kg, and there was a trend - albeit not statistically significant - for larger BC, longer IC and increased UFR and UFC when compared to non-anesthetized animals. Therefore, the use of urethane at a dose of 0.3 g/kg may be considered a good compromise between experimental/analytical ease and physiological conditions.

In order to measure bladder function in a non-invasive manner, we developed fluoroscopic volumetry, whereby subcutaneously administered contrast allowed fluoroscopic imaging of the bladder and monitoring of bladder volume during physiological filling, eliminating the need of catheter implantation. Under this condition, we found a striking, fourfold larger bladder capacity when compared to catheterized bladders undergoing (video)cystometry. In line with the findings in the catheterized animals, we observed complete voiding in awake animals and a strong reduction of the voiding efficiency after urethane anesthesia. Concurrent with the larger bladder capacities, we also measured a larger voided volume during individual voids in the non-catheterized animals, which we could attribute to longer void durations and larger peak urethral flow rates. Surprisingly, whereas urethane caused a shortening of the void duration in catheterized animals, it lead to longer voids in the non-operated animals - future research is warranted to elucidate the origin of this differential effect. The above findings indicate that the process of filling the mouse bladder via a catheter by itself has a dramatic effect on bladder function. Several mechanisms may contribute to the fourfold reduced bladder capacity. First, the implantation of a catheter into the bladder dome and fixation using a purse-string suture inevitably leads to a reduction of the area of the bladder wall and thus the bladder volume. However, since the affected surface generally does not exceed 10% of the bladder wall, this surface reduction may only explain a small part (<20%) of the capacity reduction. Second, the surgical implantation may induce local bladder irritation and inflammation, which are known to cause reduced bladder capacity (9). Lastly, filling the bladder at a supraphysiological rate of 20 µl/min, compared to the typical physiological filling rate of ∼1 µl/min in mice, may lead to increased stimulation of the bladder wall and reduced bladder capacity, as was also shown in previous research (10). These findings raise an important caveat when interpreting (video)cystometric results and highlight the usefulness of fluoroscopic volumetry to study bladder function non-invasively, albeit at the expense of intravesical pressure recordings.

In conclusion, we have combined fluoroscopy with a machine-learning-assisted analysis method to monitor the function of the LUT at high temporal resolution in mice, thereby providing a powerful approach for the non-invasive study of bladder function in awake, behaving mice.

## Materials & Methods

### Animals

Experiments were conducted on 12- to 16-weeks old female C57Bl/6J mice (Janvier). Mice were housed in filter-top cages in a conventional facility at 21 °C on a 12h light-dark cycle with unrestricted access to food and water. After each experiment, all animals were sacrificed due to the toxicity associated with urethane anesthesia. All animal experiments were carried out after approval of the Ethical Committee Laboratory Animals of the Faculty of Biomedical Sciences of the KU Leuven under project number P035/2018.

### Surgery and experimental setup

#### Urethane dose escalation

Videocystometry and suprapubic catheter implantation were performed as described earlier (3,11). In short, a PE-50 catheter was implanted into the bladder dome of five female mice through an abdominal midline incision while under 2% isoflurane anesthesia (Iso-vet, Dechra veterinary products, Bladel, The Netherlands). At the start of surgery, the anesthetized animals received carprofen (5 mg/kg bodyweight; Zoetis, Louvain-la-Neuve, Belgium) diluted in NaCl 0.9% by SC injection to achieve analgesia during videocystometry. After the animals recovered from surgery, they were placed in a small radiolucent container (4 cm x 3 cm x 7 cm) in which the animals were not restrained. Next, the container was placed in an X-ray-based small animal fluoroscopy system (LabScopeTM, Glenbrook Technologies, New Jersey, USA), and the PE-50 catheter was connected to a pressure sensor and to an infusion pump through a three-way stopcock. Videocystometry was performed with an infusion of an iodine-based intravesical contrast solution (50% Iomeron 250, Bracco Imaging Europe, Wavre, Belgium) at a rate of 20 µl/s. Simultaneously, pressure measurements were recorded at 50 Hz, and bladder images were acquired at a frame rate of 30 frames per second. Videocystometry was performed for 40 minutes, including a habituation period of 10 minutes. Next, a dose of 0.3 g/kg bodyweight urethane (Sigma-Aldrich, Diegem, Belgium) was injected subcutaneously. Ten minutes after the injection, videocystometry was performed for another 40 minutes. This procedure was repeated until a cumulative urethane dose of 1.2 g/kg was reached; at this dose, videocystometry was performed one last time, after which the experiment was terminated.

#### Catheter-free bladder imaging

Thirty minutes prior to fluoroscopy, mice were injected subcutaneously with the diuretic furosemide (15 mg/kg, 1 mg/ml solution, Sanofi, Paris, France), to increase the diuresis and thereby the filling rate of the bladder (12), and with an iodine-containing non-ionic radiocontrast agent iodixanol (12 mg I/kg bodyweight, Visipaque 320 mg I/ml; GE Healthcare, Chicago, Illinois, United States). The contrast agent was injected in the scruff area, to prevent superposition of the injected contrast solution with the bladder. Time-lapse fluoroscopy was performed on awake, freely moving mice, without bladder catheter *in situ*, during 40 minutes. Next, urethane was injected subcutaneously at a dose of 1.2 g/kg bodyweight, after which the bladder was again imaged until a void occurred.

### Data processing and analysis

As the animals were awake and freely moving, background subtraction was not possible, and the bladder volume could not be determined based on image intensity as in previous experiments (3). Therefore, we identified the bladder and traced the bladder circumference in each image, and calculated bladder volume based on the long and short radius of the identified bladder area, assuming the bladder approaches a prolate spheroid, as established in our earlier work (3):

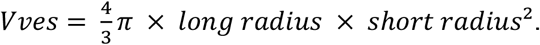

Images were saved as Audio Video Interleave file (AVI), loaded into FIJI (ImageJ) (13) and saved as an image sequence. This image sequence was loaded into NIS Elements-AR 5.30.01 (Nikon, Tokyo, Japan) and smoothed using a rolling average of 15 frames. To identify the bladder border in large sets of time-lapse fluoroscopy images, we used Segment.ai, an AI module that is a part of the NIS-Elements NIS.ai suite (Nikon, Tokyo, Japan). We trained this neural network using a set of manually annotated fluoroscopy images in which the contrast-filled bladder was imaged from various angles and at different stages of the filling/voiding cycle. The neural network was then applied to identify the bladder in similar but different datasets. It was repeatedly retrained using images where the bladder was not properly identified until satisfactory identification of the bladder border was achieved in all experiments (fig. 1a). The neural network was then used to generate an image sequence with an overlay tracing the bladder border, used for quantifying the long and short axis of the bladder and the bladder area, for large sets of time-lapse fluoroscopy images.

From the time-course of V_ves_, we determined a set of parameters related to volume alterations during urine storage and voiding (BC, RV, E_voiding_) and to urine flow during voiding (UFR, UFC), as established earlier (3), with an additional Savitzky-Golay digital filter applied to V_ves_ for data smoothing before differentiation to calculate UFR. As a measure of void duration, we determined t_20-80_, which represents the interval between the time points at which the bladder volume was reduced by 20% and 80% of the total void volume.

### Statistics

Statistical analysis and graphing was performed in OriginPro 9.0 (Originlab Corporation, Northampton, MA, USA). The Shapiro-Wilk test was used to assess normality of the data. Paired Student’s t-test, Student’s t-test and one-way repeated measures analysis of variance was used for analysis of the data. Except where indicated otherwise, all summary data are reported as mean ± SE. Animals were randomly assigned to each experimental condition. All measurements were performed on distinct animals; no animals were used in multiple experiments. Analysis was performed automatically, eliminating the need for blinding.

